# HDAC inhibitor derivatives induce differentiation of leukemic cells through two distinct and separable mechanisms

**DOI:** 10.1101/2024.01.06.572546

**Authors:** Purnima Kumar, Laia Josa-Culleré, Thomas R. Jackson, Carole J. R. Bataille, Paresh Vyas, Thomas A. Milne, Angela J. Russell

## Abstract

Acute myeloid leukaemia (AML) is a haematopoietic malignancy comprising different genetic subtypes with a common hallmark of differentiation arrest. In abnormal haematopoiesis, overcoming the differentiation blockade has emerged as an attractive therapeutic strategy. In a screen with genetically distinct AML cell lines, histone deacetylase inhibitors (HDACis) were observed to cause an upregulation in the expression of CD11b, a myeloid differentiation marker. These caused changes in cell morphology, block in proliferation, and cell cycle arrest at the G1 phase. To gain insights onto the mechanism of these compounds, we planned to prepare inactive probes devoid of the zinc binding motif. However, these compounds were unexpectedly still able to initiate differentiation, albeit through a distinct target and via a G2 arrest. Subsequent RNA sequencing studies supported the differentiation phenotype for the HDACis and highlighted the role of cell cycle regulatory kinases for the effect observed in the probe molecules. We then showed that these inhibit Aurora A and GSK3α kinases, suggesting their potential as therapeutic targets for differentiation therapy in AML. Our work supports the importance of properly validating inactive tool compounds and their potential to identify novel targets.

## Introduction

Acute myeloid leukaemia (AML) is a disease of the blood and the bone marrow. It is a well-characterised haematological malignancy that is marked by uncontrolled proliferation of leukemic cells, decreased apoptosis, and a block in differentiation, leading to an increase of immature cells.^1^ These immature cells accumulate in the bone marrow affecting the formation of other blood cell lineages, thereby interfering with normal haematopoiesis.

The inhibited differentiation in AML cells is caused by both genetic abnormalities and epigenetic changes such as DNA hypermethylation and aberrant histone acetylation.^2^ Recently, overcoming the differentiation block has been proposed as an attractive approach that could overcome the disadvantages of current treatments, such as high toxicities from chemotherapy and the development of resistance from “targeted therapy”.^3^ The first example of such differentiation approach was the use of all-trans retinoic acid (ATRA), which showed granulocytic differentiation of HL60 leukemic cells^4^ and was later approved by the FDA in 1995 for the treatment of acute promyelocytic leukaemia (APL). This massively improved the survival rates of patients with this sub-type of AML.^5^

In spite of the huge impact of differentiation therapy with ATRA, this approach is based on targeting an oncoprotein that is specific for patients with APL, thus it is not effective for the other 85-90% of patients.^6^ Therefore, there is a need to find alternative targets that can be exploited to identify new differentiation agents that are effective in wider patient populations. In this context, phenotypic screening is a promising approach that permits identifying small molecules with new mechanisms of action. In fact, an analysis of the drugs approved for AML revealed that the largest proportion originates from phenotypic drug discovery (PDD) campaigns.^7^ However, when we started this work there was no precedent on the use of phenotypic screens based on an AML cell differentiation readout.

We recently described how we set up a phenotypic screening platform to identify small molecules that could differentiate AML cells of various sub-types.^8^ We used this platform to identify hits from several chemical series.^9–11^ Our primary assay was based on detecting the expression of CD11b, which is overexpressed in differentiated myeloid cells, by flow cytometry after a 4-day treatment of HL60 cells. In order to validate this assay, we had previously performed a small pilot screen of 90 small molecules with known targets. Amongst these, several histone deacetylase (HDAC) inhibitors, such as mocetinostat and an aliphatic derivative **1** (Figure 1A), showed a robust ability to differentiate HL60 cells. Herein, we present our follow-up work on these HDACis.

**Figure 1.**
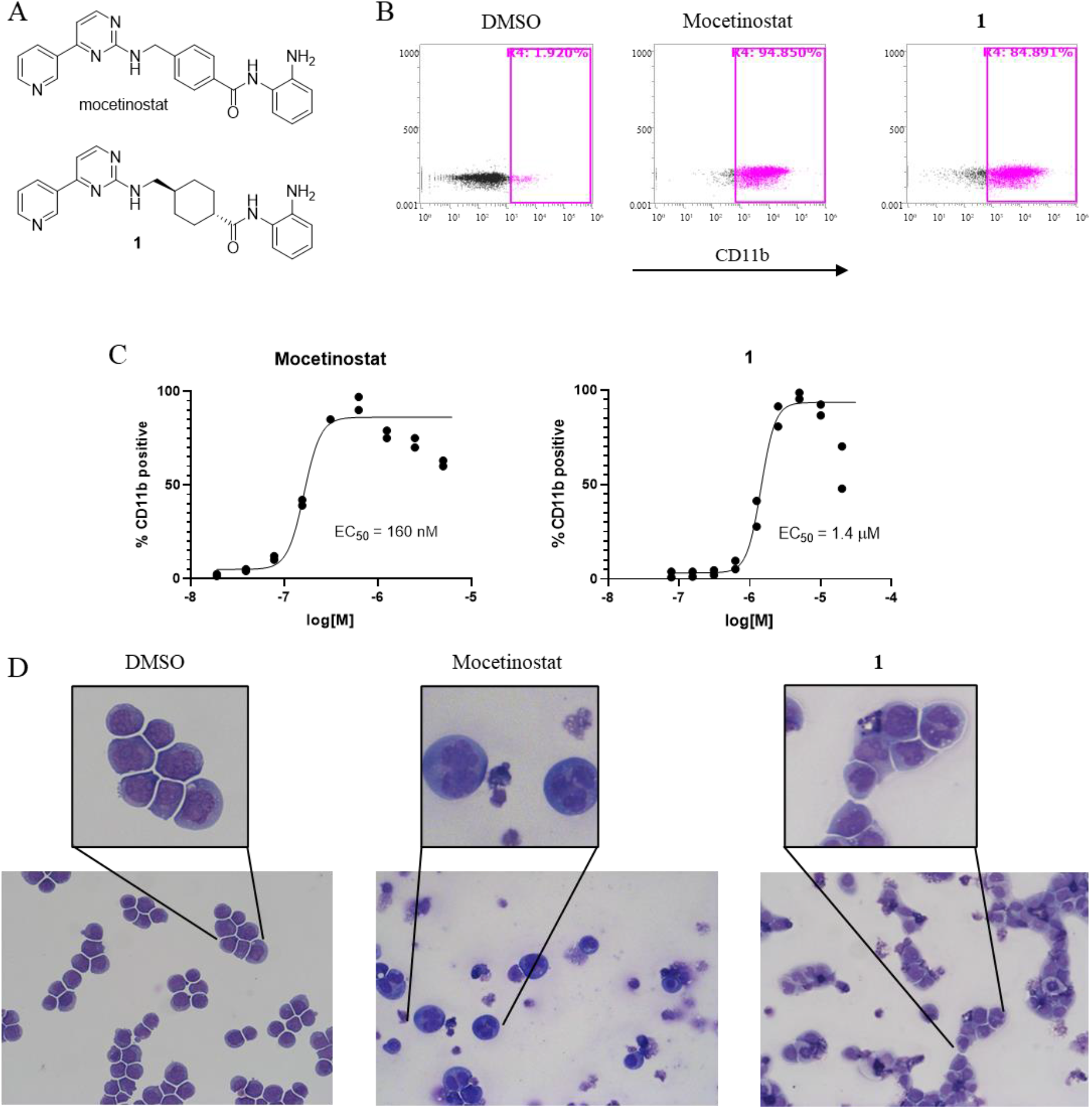
Mocetinostat and compound **1** induce differentiation in AML cells. (A) Chemical structures for mocetinostat and **1**. (B) Flow cytometry plots of CD11b expression of HL60 cells treated with either DMSO control, mocetinostat (300 nM) or **1** (5 μM) for 4 days. (C) Representative concentration-response plots of HL60 cells treated for 4 days with mocetinostat (EC_50_ = 160 nM) and **1** (EC_50_ = 1.4 μM). (D) Cytospins of HL60 cells stained with Wright-Giemsa showed morphologic changes such as presence of vacuoles, multiple nucleoli and large cytoplasm when treated with mocetinostat (300 nM) or **1** (2.5 μM) compared to DMSO control cells.

Histone deacetylases (HDACs) have been implicated in multiple biological processes, most famously regulating the balance of acetylation on chromatin thus influencing gene regulation.^12^ However, HDACs can also impact processes as diverse as metabolism and the maintenance of formaldehyde levels in the cell.^13^ For this reason, small molecule HDAC inhibitors (HDACis) have been found to have a wide range of effects including impacts on cell differentiation, apoptosis, as well as growth arrest.^14^ Short-chain fatty acids such as 4-phenylbutyrate and valproic acid have been explored to study the effects of HDACis both *in vitro* and in clinical trials.^15^ However, one of the concerns raised with this class of compounds is their off-target effects limiting their specificity as well as therapeutic potential.^16^ HDACis containing hydroxamic acids (such as SAHA, pyroxamide, TSA, oxamflatin, and CHAPs) have been postulated to interact with the catalytic zinc binding domain via coordination of the hydroxamate moiety in a bidentate fashion, thereby blocking substrate access, as shown by X-ray crystallographic studies.^17,18^ Another important class of HDACis are the ones having a benzamide group, which interact with the catalytic site in a similar manner binding to the active zinc atom. This class of HDACi includes compounds such as mocetinostat and MS-275, which have been studied for AML and are currently used in clinical trials in combination with other cytotoxic therapies.^19^ MS-275 has been shown to exert a dose-dependent effect on the AML cell line U937, with a p21-dependent growth arrest causing differentiation at low concentrations, followed by apoptosis at higher concentrations.^20^

The mechanism by which the HDACi mocetinostat induces differentiation of AML cells is not fully understood. To gain some insight, we performed morphology, cell cycle, and RNA sequencing studies to deconvolute the downstream signalling pathways that lead to differentiation, which we present herein. We planned to support these studies by preparing some presumably inactive probes where the *ortho*-amino anilide motif, which constitutes the key zinc binding group, is altered. However, we found these compounds to still cause differentiation of our AML cell lines. We also present some mechanistic characterisation of these compounds as well as potential new targets for differentiation.

## Results

### Validation of hits

CD11b is a known marker of myeloid differentiation and was used to monitor the induction of differentiation of AML cell lines. A phenotypic screen was performed with an in-house library of compounds that included some with known targets (data not shown). From this screen, the well-known HDACi mocetinostat and its novel saturated analogue **1** (Figure 1A, Scheme S1) were obtained as hits, increasing CD11b expression up to 85-95% (Figure 1B). In order to determine the potency of these compounds in three AML cell lines HL60, THP1 and OCI-AML3, cells were treated with various concentrations of the compounds for 4 days. Cells were then stained with the differentiation marker CD11b, and its expression was determined by flow cytometry. Mocetinostat and **1** had EC_50_ values of 160 nM and 1.4 μM respectively in HL60 cells (Figure 1C), 1.5 μM and 1.8 μM in THP1 cells (Figure S1), and 5.2 μM and 6 μM in OCI-AML3 cells (Figure S2). To further validate the hits, morphology of the cells was characterised using Giemsa staining, which showed clear changes upon compound treatment, such as the presence of vacuoles, multiple nucleoli, and an extended cytoplasm as compared to the DMSO treated cells, consistent with myeloid differentiation (Figure 1D). These HDAC inhibitors thus proved to induce differentiation in a range of cell lines with different genetic characteristics.

We were interested in understanding how HDAC inhibition induces differentiation of the AML cells. For this purpose, we envisioned that preparing close analogues of the validated hits devoid of the zinc-binding motif could serve as mechanistic probes. It is known that the *ortho*-aminobenzamide moiety of these compounds is essential for HDAC inhibition, as it can chelate the zinc atom on the active site,^21^ hence the *meta* analogues **2** and **3** (Figure 2A) were prepared (Scheme S2) as potential negative controls. To our surprise, these compounds were still able to induce differentiation of the AML cells, albeit at lower potency (Figure 2B, 2C, and S3). At 40 μM, these increased %CD11b up to 60%. The differentiation effect was also further confirmed by morphology analysis (Figure 2D). A Western blot analysis looking at histone 3 acetylation after 24 h of treatment with the compounds confirmed that **1**, but not its corresponding *meta* analogue **3**, increased acetylated levels (Figure 2E and 2F).

**Figure 2.**
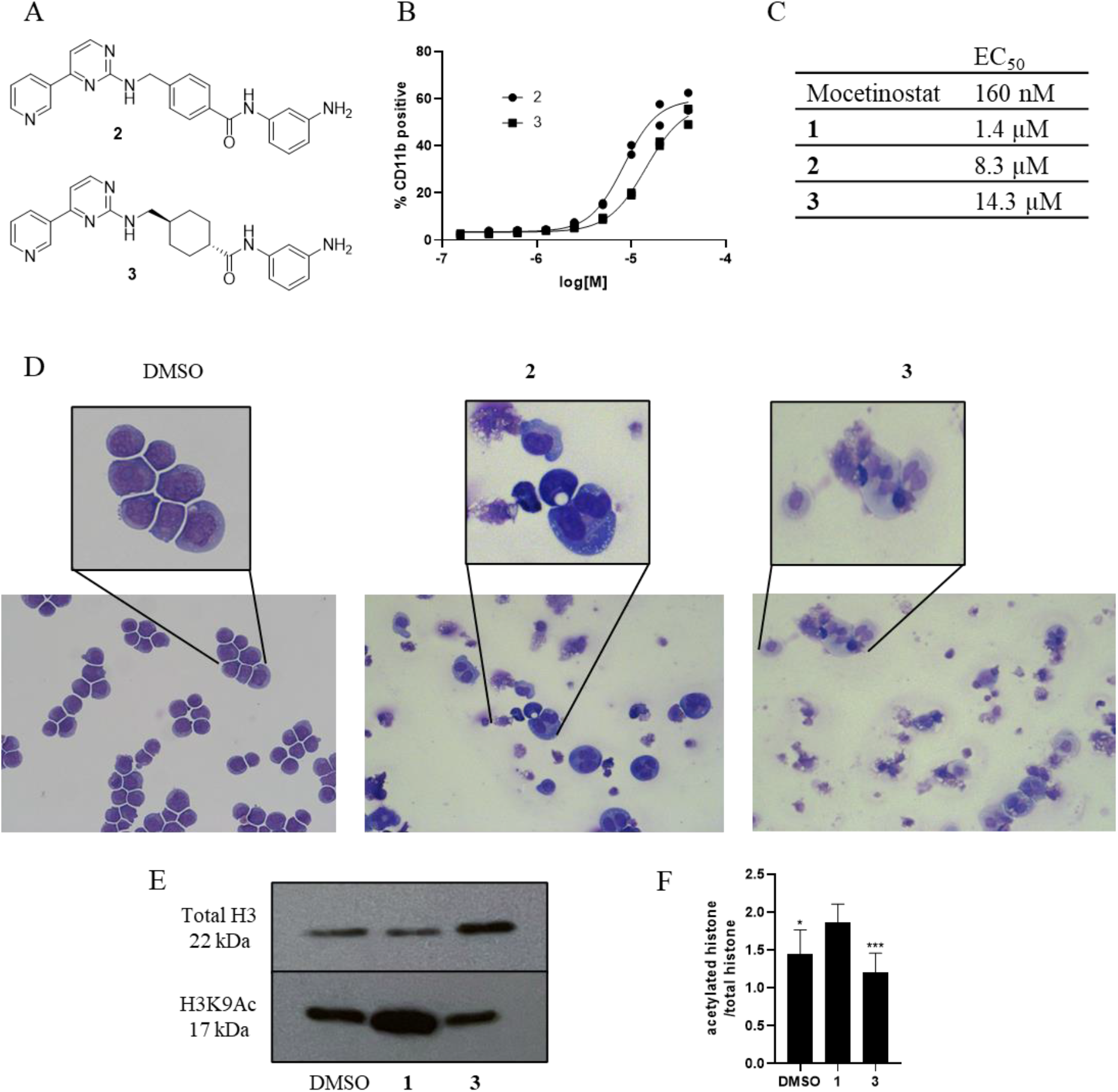
Compounds **2** and **3** induce differentiation in AML cells but are not HDACis. (A) Chemical structures of **2** and **3**. (B) Concentration-response of for HL60 cells treated for 4 days with **2** and **3**. (C) EC_50_ values of the *ortho* and *meta* analogues for HL60 cells treated for 4 days with the indicated compounds. (D) Cytospins of HL60 cells stained with Wright-Giemsa showing morphologic changes when treated with **2** or **3** (30 μM) compared to DMSO control cells. (E) Western blot analysis of the effect of **1** (2.5 μM) and its *meta* analogue **3** (30 μM) on histone H3 acetylation after 24 h treatment in HL60 cells. (F) Quantification of acetylated histone 3 level normalised to total histone 3 with treatment of **1** and **3**, optical intensity measured with ImageJ. Error bars shown as SEM of n=3. P values were calculated using One-way ANOVA. *P<0.05, ***P< 0.001.

Recombinant enzyme assays were used to confirm HDAC inhibition activity and selectivity of the compounds. The compounds were tested against different HDACs belonging to different groups, all of which are zinc-dependent enzymes. It was confirmed that HDAC1, 2, and 11 were inhibited by mocetinostat,^22^ while **1** inhibited only HDAC1 and 11 (Table 1). Also, the *meta* analogues **2** and **3** failed to inhibit any of the four HDACs, and thus were confirmed as non-HDACis, implying that the phenotypic effects observed with the *meta* analogues originate from an alternative mechanism of action.

**Table 1.**
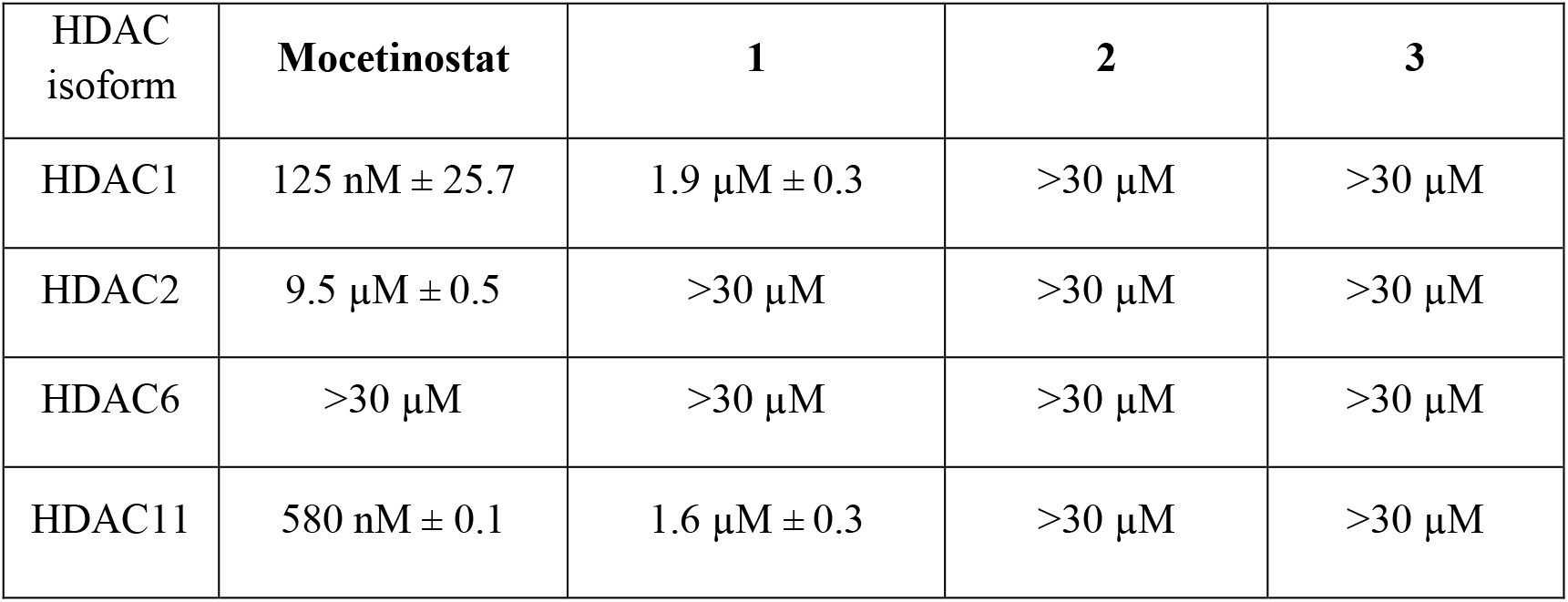
IC_50_ values of the *ortho* and *meta* analogues against recombinant human HDACs. Curves shown in Figure S4. Results are calculated as means ± SD based on experiments performed in biological triplicate (n=3).

Due to the dysregulation of proliferative factors, leukemic cells proliferate in an uncontrolled manner. However, upon inducing differentiation, a block in proliferation is usually observed. To confirm this for our compounds, cells were stained with acridine orange and DAPI to measure live cell number and viability after 1 – 4 days of treatment. The four compounds induced a visible reduction in the cell number as compared to the negative control in the three cell lines HL60 (Figure 3A and 3B), THP1, and OCI-AML3 (Figure S1C and S2C). A timepoint study of CD11b expression was also performed where the cells were treated with the respective compounds for 4 – 96 h. While upregulation of CD11b was already detected after 24 h with both *ortho* and *meta* derivatives (Figure 3C), the effect on proliferation is observed more strongly after 72 h, consistent with the proliferation block being a downstream effect of differentiation. Also, while the *meta* analogue **2** decreased cell numbers to the same extent than the corresponding *ortho* compound, it did not have a pronounced effect on cell viability after 4 days.

**Figure 3.**
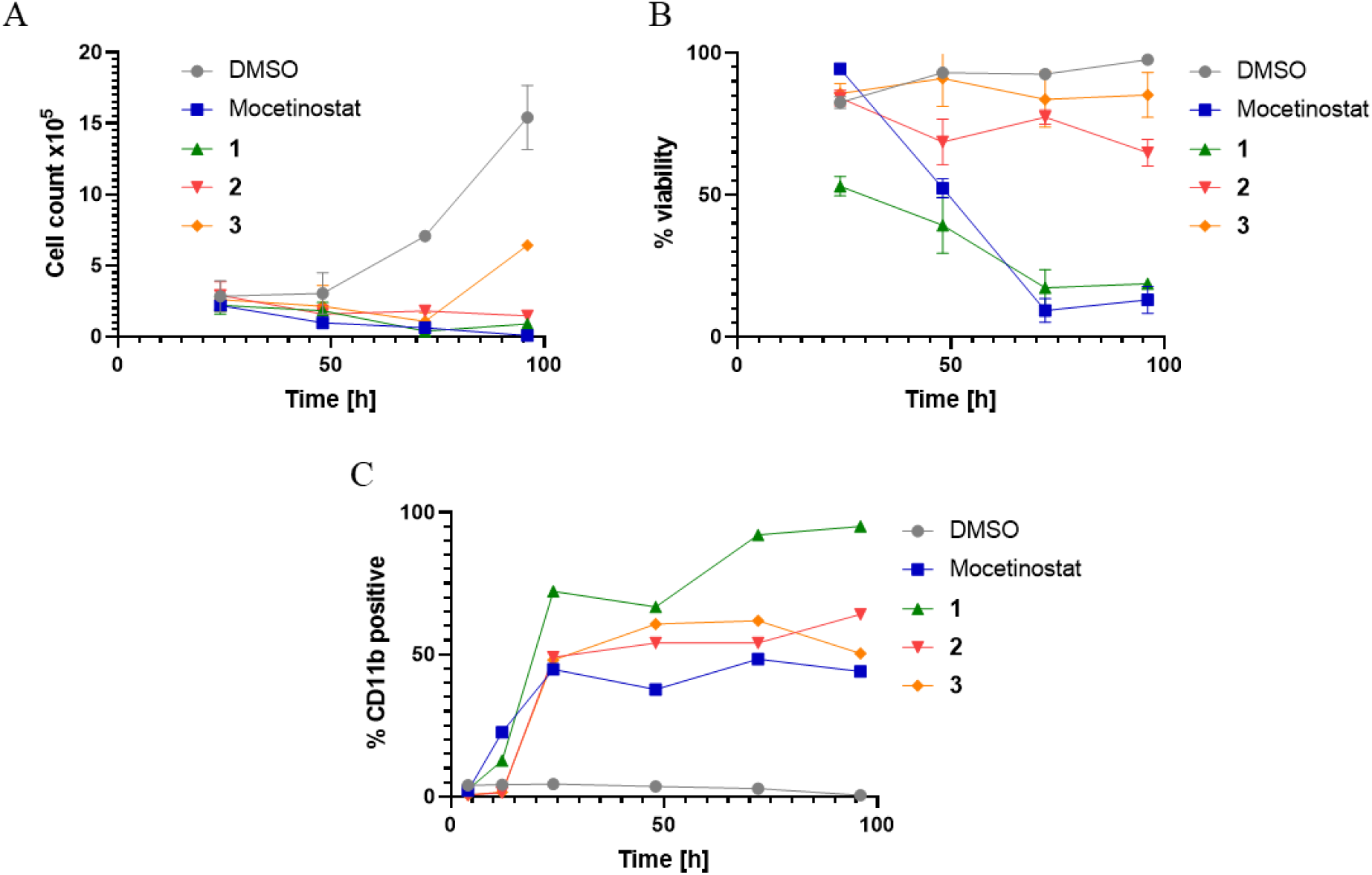
O*rtho* and *meta* analogues inhibit cell proliferation in HL60 cells. HL60 cells were treated with the compounds (600 nM mocetinostat, 2.5 μM **1**, 30 μM **2** and 30 μM **3**) and (A) live cells per well and (B) cell viability were measured via acridine orange and PI staining, and (C) CD11b expression by flow cytometry at the indicated timepoints. Error bars shown as SEM of n=3.

### Cell cycle analysis

In a regular cell cycle, cells can exit the G1 phase and enter a state of quiescence (G0), from where they can either undergo differentiation or re-enter cell cycle to proliferate.^23^ In AML, mutated pathways play a role in the regulation of cell division. Given the differentiation signature observed for the compounds, an arrest on G1 could be expected. To explore this in detail, we examined the effect of the HDACis mocetinostat and **1** on cell cycle and compared it with that of the *meta* analogues. HL60 cells were treated with the compounds for 48 h, and then stained with PI and analysed by flow cytometry. The results confirmed that the *ortho* analogues caused a G1 arrest, however the *meta* analogues induced an increase of G2 instead (Figure 4). The data was consistent for OCI-AML3 and THP1 cells (Figure S5). The remarked change in the effect of this compounds on cell cycle depending on such a small chemical modification is noteworthy.

**Figure 4.**
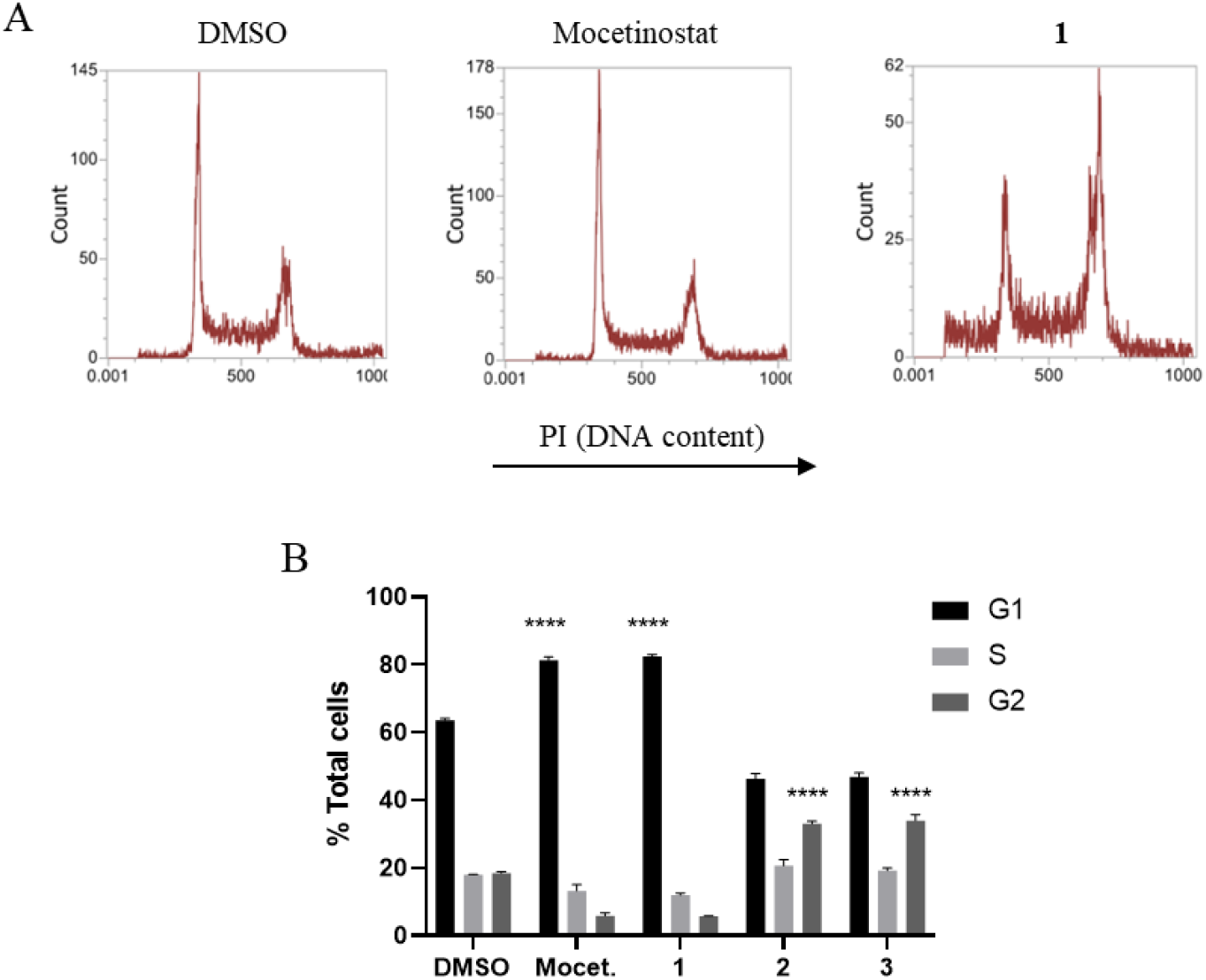
The *ortho* compounds cause a G1 arrest and the *meta* analogues a G2 arrest. HL60 cells were treated with the compounds (600 nM mocetinostat, 2.5 μM **1**, 30 μM **2** and **3**) for 48 h, stained with PI, and analysed by flow cytometry. (A) Representative flow cytometry plots. (B) Quantification of %cells in G1, S, and G2 phases of the cell cycle. Error bars shown as SEM of n=3. P values were calculated using Two-way ANOVA, ****P< 0.0001.

### RNA sequencing

An RNA sequencing study was carried out to identify any significant changes in RNA expression resulting from compound treatment. The samples were sequenced using poly-A RNA sequencing. HL60 cells were treated with the respective compounds for a period of 24 h followed by RNA extraction. DMSO (0.1%) samples were treated as the negative control. The Volcano plots of mocetinostat and **1** showed a significant difference (P_adj_<0.05) in the regulation of gene expression as compared to the negative controls (Figure 5A). There was a lower number of differentially regulated genes between the *meta* analogues and the negative controls. The results from the heatmap in Figure 5B also indicated that **1** and mocetinostat were clustered further from the negative controls, highlighting difference between their gene regulation patterns, while the *meta* analogues clustered closely with the DMSO samples. Mocetinostat and **1** looked similar between themselves at global gene expression level, with few differences in up-regulated and down-regulated genes (Figure 5C).

**Figure 5.**
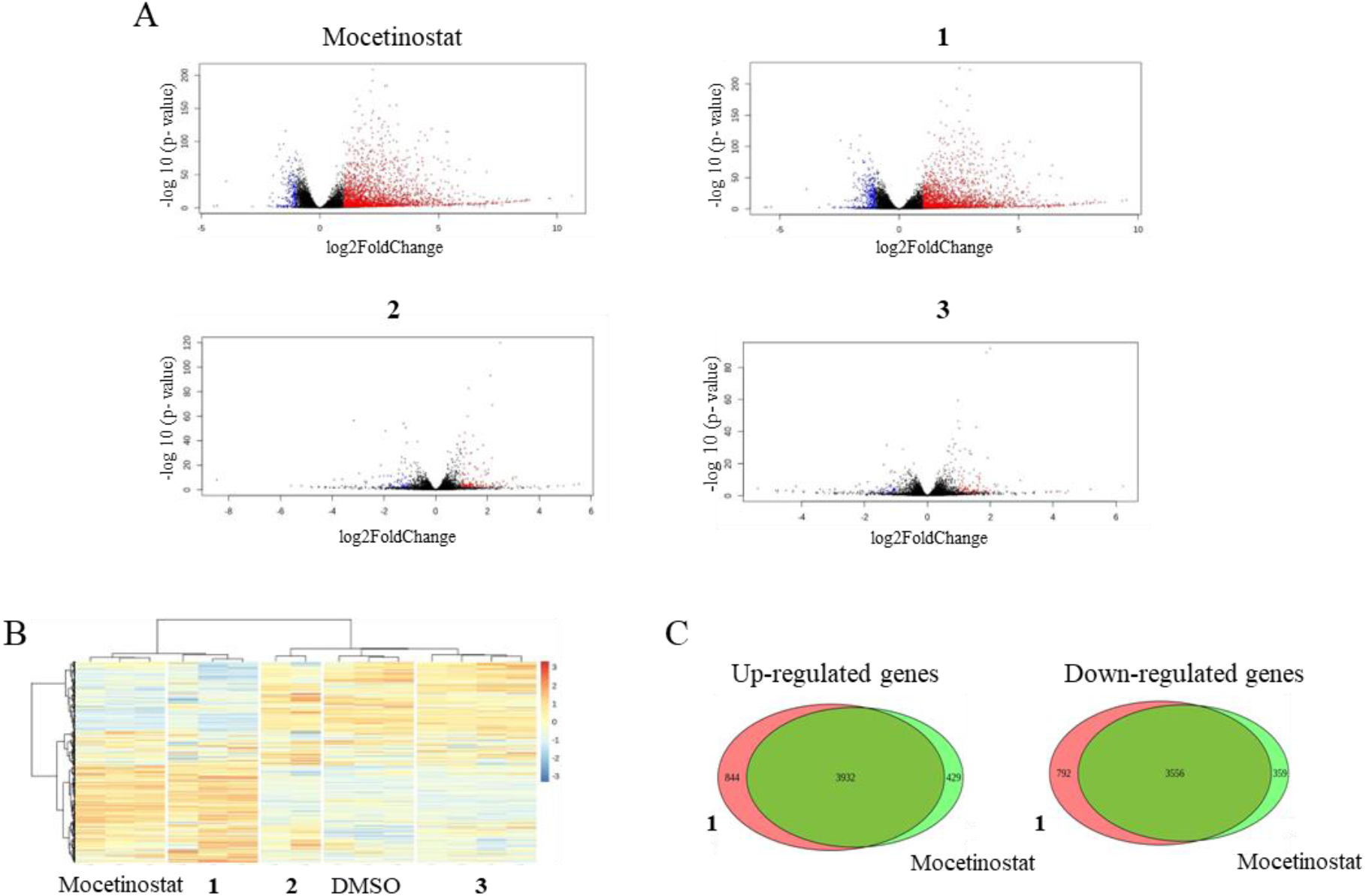
RNA sequencing reveals gene expression changes of HL60 cells upon 24 h treatment with compounds (600 nM mocetinostat, 2.5 μM **1**, 30 μM **2** and **3**). (A) Volcano plot of the differentially expressed genes between normal vs compound treated cells. Significantly down-regulated genes are in dark blue, significantly up-regulated genes are in red, non-significant genes are in black. (B) Heatmap of hierarchical clustering indicating differentially expressed genes (rows) between DMSO control and compounds (fold-change > 1, P < 0.05). Orange indicates upregulation and blue indicates downregulation. (C) Venn diagram of differentially expressed genes between cells treated with either mocetinostat or **1**.

EnrichR analysis revealed that **1** and mocetinostat up-regulated genes which overlapped with macrophage and neutrophil signatures (Figure S6 and S7), confirming the differentiation phenotype previously characterised. As compared to gene expression changes of other molecules in the L1000CDS2 database, the top-ranking matches were for known HDAC inhibitors. Conversely, the gene signatures for *meta* analogues **2** and **3** closely matched with a few kinase inhibitors (Figure S8). To confirm these results, the compounds were tested in a kinase inhibitory screen. The *meta* analogues, but not the *ortho*, were found to inhibit Aurora-A kinase and GSK3 alpha kinases (Figure 6).

**Figure 6.**
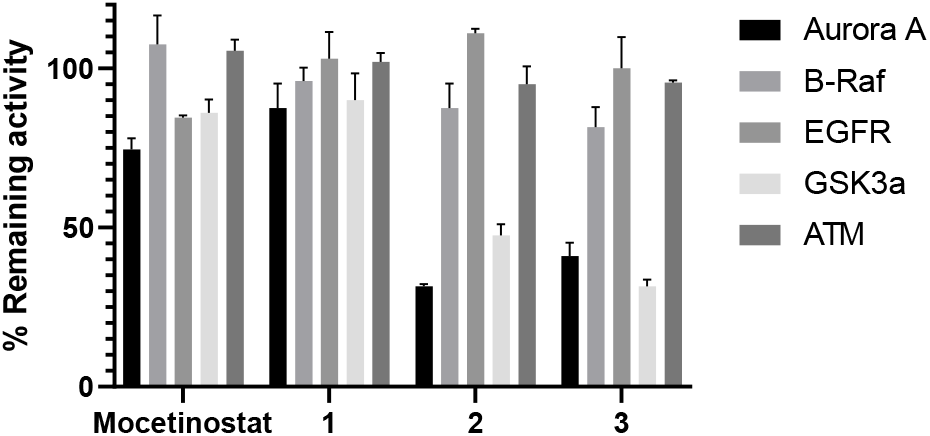
A kinase screen identifies that the *meta* analogues **2** and **3** inhibit Aurora A and GSK3α. Remaining kinase activity was measured after treatment with the compounds (600 nM mocetinostat, 2.5 μM **1**, 30 μM **2** and **3**). Error bars shown as SEM of n=2.

## Discussion

HDAC inhibitors such as panobinostat, vorinostat, and TSA promote cell death, autophagy, and apoptosis in preclinical AML models and are also used at various stages of clinical development, yet these compounds are not effective as monotherapy primarily due to dose-limiting tolerance attributed to off-target effects.^19^ Hence, there is a need to develop more specific HDAC inhibitors with improved profile and therapeutic window. A greater understanding of the underlying mechanism and off-target effects caused by HDAC inhibition would in turn aid in the design of isoform-selective HDAC inhibitors.

In this study, a screen of compounds with different targets revealed that HDACis cause differentiation in a variety of genetically distinct AML cell lines. Besides mocetinostat, a saturated analogue **1** showed a similar profile. As a higher number of aromatic rings in drugs correlates with poorer compound developability as an oral drug and increased risk of attrition,^24^ compound **1** represents an interesting addition to the toolbox of HDAC inhibitors. Previous studies have shown that the *ortho*-aminoanilide moiety binds potently to the active zinc ion via bidentate coordination at the catalytic site, leading to the HDAC inhibitory effects.^17,18^ To confirm if HDAC inhibition leads to differentiation, potential inactive analogues were prepared by shifting the amino group to the *meta* position. However, the resulting compounds **2** and **3** still caused an upregulation of CD11b expression. These results suggested that these new compounds were giving rise to the differentiation phenotype through an alternative mechanism.

An analysis of the effects of the compounds on cell cycle revealed that the *ortho* analogues caused a G1 arrest, while the *meta* analogues caused a G2 arrest, further confirming the distinct mechanism of the compounds. These results show that a simple change in the structure of small molecules can lead to a prominent change on their cellular effect; in this case, switching from a G1 to a G2 arrest in the cell cycle. Further, RNA sequencing data supported the differentiation signature of the *ortho* analogues, and it also suggested kinase inhibition as the underlying mode of action of the *meta* analogues. A kinase screen then identified that **2** and **3** inhibit the kinases Aurora A and GSK3α at the same concentration at which they generate a cellular effect. These results support previous work where GSK3α was identified via a genetic screen as a target in AML.^25^ It is also consistent with recent work showing that inhibition of Aurora A and B causes a G2 arrest in t(8;21) AML cells,^26^ that a novel Aurora A inhibitor PW21 inhibits proliferation of leukemic cells by attenuating their interaction with the mesenchymal stem cell microenvironment,^27^ and that the Aurora A inhibitor alisertib induces differentiation of THP1 cells through the KDM6B pathway.^28^

Phenotypic drug discovery (PDD), which does not rely on prior knowledge of the identity of a specific drug target and a hypothesis about its role in disease,^29^ seems a truly promising approach to identify small molecules that can differentiate AML cells regardless of their mutation status. However, despite an increasing interest in PDD in the recent years, this strategy does come with challenges, especially in the hit validation and target deconvolution phases. In fact, the majority of successful drug discovery programmes seem to combine target knowledge and functional cellular assays to identify drug candidates with the most advantageous molecular mechanism of action (MoA).^30,31^ Gaining insight on the target and MoA of small molecules often relies on high-quality tool compounds, which can support the challenging task of distinguishing between on-target and off-target effects. However, identifying suitable tool compounds is a challenging task, and indeed a systematic literature review revealed suboptimal use of chemical probes in cell-based biomedical research.^32^ To control for any off-target activity of the developed molecules, an inactive close analogue is often used as a negative control.^33^ This is often based on an inactive enantiomer for chiral molecules or on a small modification that completely abolishes the phenotypic effect.

All in all, in this work, we demonstrate how we initiated mechanistic studies with seemingly highly specific compounds targeting HDACs. To obtain inactive probes that would support these studies, we modified them by abolish their ability to bind to the zinc atom in the HDAC active site. While this consistently eliminated HDAC activity, these “inactive” probes were unexpectedly still able to differentiate AML cells. Follow-up RNAseq studies pointed to kinases as possible drivers of the phenotypic effect. Ironically, our previously published phenotypic screen, which was completely target-agnostic, and follow-up MoA studies turned out to identify compounds that bind to tubulin, which is a well-known target in oncology,^8^ albeit previously unknown for AML differentiation. In contrast, in the work described in this publication we started with a known mechanism and ended up with a completely different target for differentiation therapy.

Our work represents a cautionary tale on the design of inactive probes, on the importance to properly validate chemical tools, and on how a small chemical change can lead to completely different cellular modes of action. It also supports the power of RNA sequencing as a key approach for target deconvolution strategies. Our compounds **1**-**3** also represents an exciting addition to the literature by expanding the available toolbox of differentiation agents in AML; these are not only effective in a particular patient population like other targeted therapies, but instead work across cell lines with different genetic characteristics. This work also supports Aurora A and GSK3α as promising targets to identify compounds that can induce G2 cell cycle arrest and differentiation to AML cells.

## Methods

### Compounds

10 mM stock solutions were prepared in DMSO and stored at -20 ºC. Serial dilutions of the stock solutions were carried out prior to use in each experiment and final concentration of DMSO was maintained at 0.1% except for final compound concentrations above 10 μM. Non-commercial compounds were synthesised as described in the Supporting Information.

### Cell culture

Human leukaemia cell lines THP1, HL60 and OCI-AML3 were supplied by ATCC. All cell culture reagents were purchased from Invitrogen Life Technologies and cells were maintained in a humidified incubator set to 37 ºC, 5% CO_2._ Cells were grown in RPMI containing 10% FBS and 1% L-Glutamine.

### Reagents

PE mouse anti-human CD11b (clone ICRF44) and IgG isotype (clone MOPC-21) were purchased from BD Biosciences and DAPI from Sigma. Anti-rabbit acetylated H3 was purchased from Millipore. Antirabbit H3 and anti-rabbit IgG were purchased from Abcam Plc. Ph3 antibody Alexa-Fluor 488 anti-histone H3 phospho (Ser10) (clone 11D8) was purchased from Cell Signalling Technology.

HDAC inhibition recombinant assay enzyme kit was purchased from Enzo Life Sciences and was performed according to the manufacturer’s protocol.

### Flow cytometry antibody staining

Exponentially growing cells were seeded in a 96 well plate at a density of 2x10^5^ cells/mL in the presence of the respective compounds in a final volume of 100 μL. The plates were incubated for 4 days. Cells were pelleted by centrifugation at 1000 rpm and pellets were resuspended in 40 μL of blocking buffer (10% FBS in IMDM) containing either anti-CD11b or IgG isotype. The cells were stored in ice for 20 min. Cell suspension was then centrifuged, washed three times with staining buffer (1% FBS in IMDM), and resuspended in 200 μL of staining buffer with 1 μg/mL DAPI. The cells were analysed through Attune NxT flow cytometer (Thermo Fisher Scientific) with previous compensation. Data was analysed using Invitrogen Attune NxT software (Thermo Fisher Scientific) and Flow Jo (v9).

### Wright Giemsa staining

Cells were prepared at approximately 2x10^5^ cells/ml in 200 μL of staining buffer and loaded into cytospins at 1000 rpm for 10 min. Cells were stained with Modified Wright’s stain using a Hematek 200. The slides were then covered with a glass cover slip and viewed under Olympus BX-30 and analysed with the Infinity software.

### Cell proliferation and viability assay

Exponentially growing cells were plated in a 96 well plate at a density of 2x10^5^ cells/mL in a final volume of 100 μL. Cells were incubated in the presence of 10 μM and 1 μM of the respective compounds for 1 – 4 days. Viable cell number was determined using Solution 13 (ChemoMetec) containing acridine orange and DAPI and analysed using the Nucleo Counter NC-3000 system (ChemoMetec) according to manufacturer’s protocol. Results were analysed using Graphpad Prism Software.

### Cell cycle analysis with flow cytometry

Exponentially growing cells were plated in a 96 well plate at a density of 2x10^5^ cells/mL. Cells were incubated with the respective compounds and their respective concentrations in a final volume of 100 μL for 1 – 3 days. Cells were pelleted by centrifugation at 600 x g for 6 min at room temperature. Pellets were washed once with PBS and stained with propidium iodide solution (0.1% v/v Triton X 100, 50 μg/mL propidium iodide, 200 μg/mL RNAase in PBS) for 2 h on ice. Samples were analysed by Attune NxT flow cytometer and analysed with Invitrogen Attune NxT software.

### RNA sequencing

5x10^7^ cells/mL were used with a total of 5x10^6^ cells per sample. Cells were treated with the compounds (600 nM mocetinostat, 2.5 μM **1**, 30 μM **2** and 30 μM **3**) for 24 h. RNA was extracted using QIAGEN RNeasy-Plus Mini columns as per the manufacturer’s instructions. RNA purity was analysed using RNA Screen Tape with a TapeStation system (Agilent). The samples were sent to Oxford Genomics Centre, Welcome Centre for Human Genetics for poly-A RNA sequencing.

All data represent at least three independent experiments and were expressed as mean ± standard error of the mean (SEM). Statistical analyses were performed using Student’s *t*-test for comparison between two groups, and analysis of variance (ANOVA) and Dunnett’s post hoc test for multiple comparisons among groups. A probability value of *p* < 0.05 was considered significant. GraphPad Prism 5 software version 5.01 was used for data analyses.

## Supporting information

Supplementary Figures and Schemes, Chemistry Experimental Procedures

## Data availability

The data presented in this study is contained within the article and supplementary material. The datasets are available from the corresponding author on reasonable request.

## Acknowledgements

T.A.M. is supported by Medical Research Council (MRC, UK) Molecular Haematology Unit grant MC_UU_00016/6. T.R.J., L.J-C., and the experimental data were all supported with a grant from OxStem Oncology.

## Author contributions

P.K., L.J-C., T.R.J., C.J.R.B., P.V., A.J.R. and T.A.M. conceived the experimental design; P.K., L.J-C., T.R.J., and C.J.R.B. carried out experiments, analysed, and curated the data; K., L.J-C., T.R.J., C.J.R.B., A.J.R. and T.A.M. interpreted the data; P.K., L.J-C., A.J.R. and T.A.M wrote the manuscript; all authors contributed to reviewing and editing the manuscript; P.V., A.J.R. and T.A.M. provided supervision and funding.

