## Supplementary Figures and Schemes, Chemistry Experimental Procedures for "HDAC inhibitor derivatives induce differentiation of leukemic cells through two distinct and separable mechanisms"

### Table of contents

**Supplementary Figures and Schemes** ........................................................................ S3

**Figure S1**: Mocetinostat and **1** induce differentiation in THP1 cells.

**Figure S2**: Mocetinostat and **1** induce differentiation in OCI-AML3 cells.

**Figure S3**. Flow cytometry plots of CD11b expression of HL60 cells.

**Figure S4**: Dose-response curves of compounds in HDAC recombinant assays.

**Figure S5**. Effect of compounds on cell cycle.

**Figure S6**. EnrichR analysis of mocetinostat.

**Figure S7**. EnrichR analysis of **1**.

**Figure S8**. EnrichR analysis of **2** and **3**.

**Figure S9**. Western blot of **1** and **3**.

**Scheme S1**. Synthesis of mocetinostat and its *meta* analogue **2**.

**Scheme S2**. Synthesis of **1** and its *meta* analogue **3**.

**Chemistry Experimental Procedures** ........................................................................ S11

**References** ........................................................................................................ S16

### Supplementary Figures

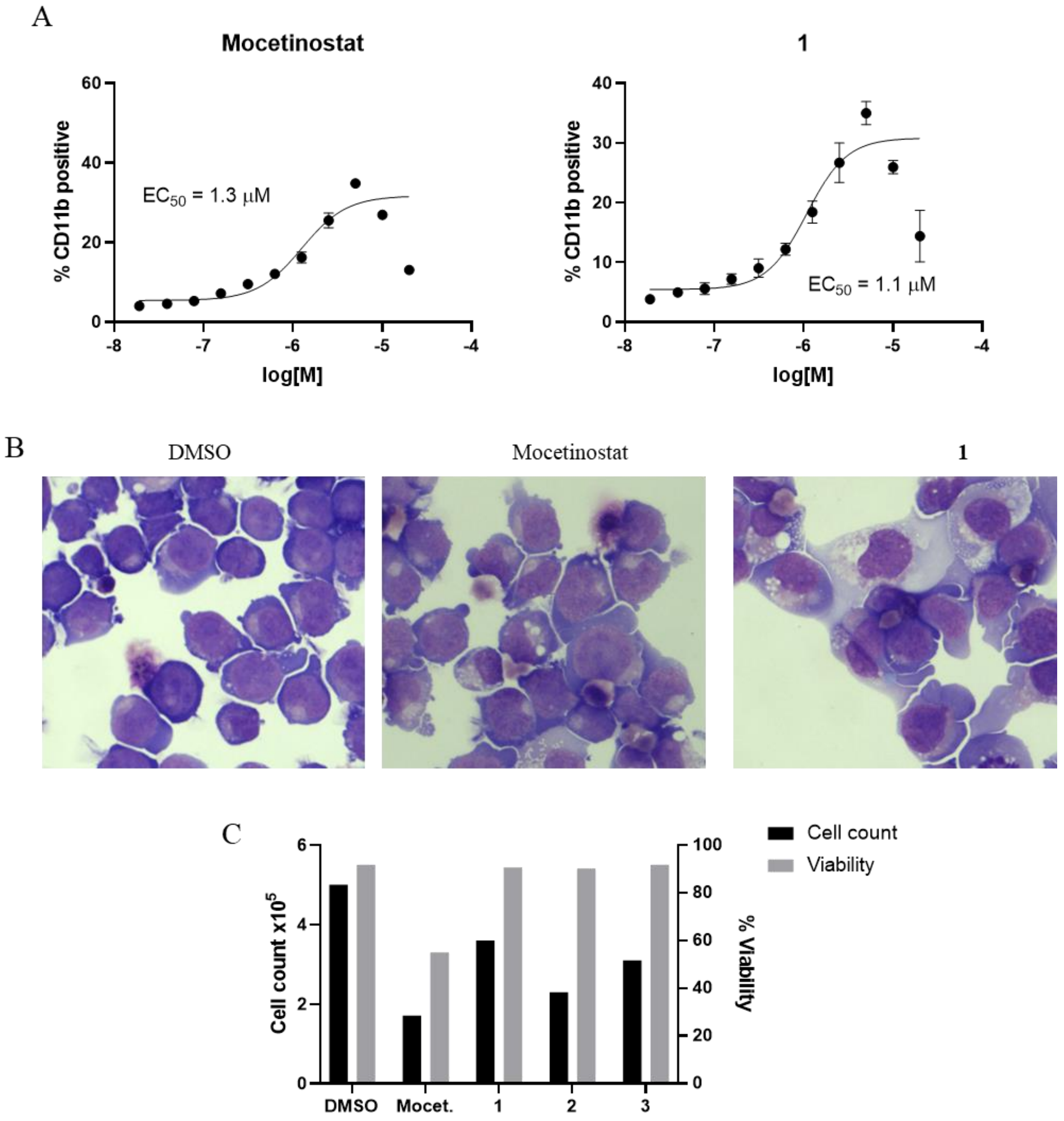


**Figure S1**: Mocetinostat and **1** induce differentiation in THP1 cells. (A) Concentration-response curves of %CD11b expression in THP1 cells treated with mocetinostat and **1** for 24 h, measured by flow cytometry. (B) Cytospins of THP1 cells stained with Wright-Giemsa showed morphologic changes consistent with differentiation when treated with mocetinostat (600 nM) or **1** (2.5 µM) compared to DMSO control cells. (C) Live cells per well and cell viability measured via acridine orange and PI staining of THP1 cells treated with the compounds (600 nM mocetinostat, 10 µM **1**, 80 µM **2** and **3**) for 4 days.

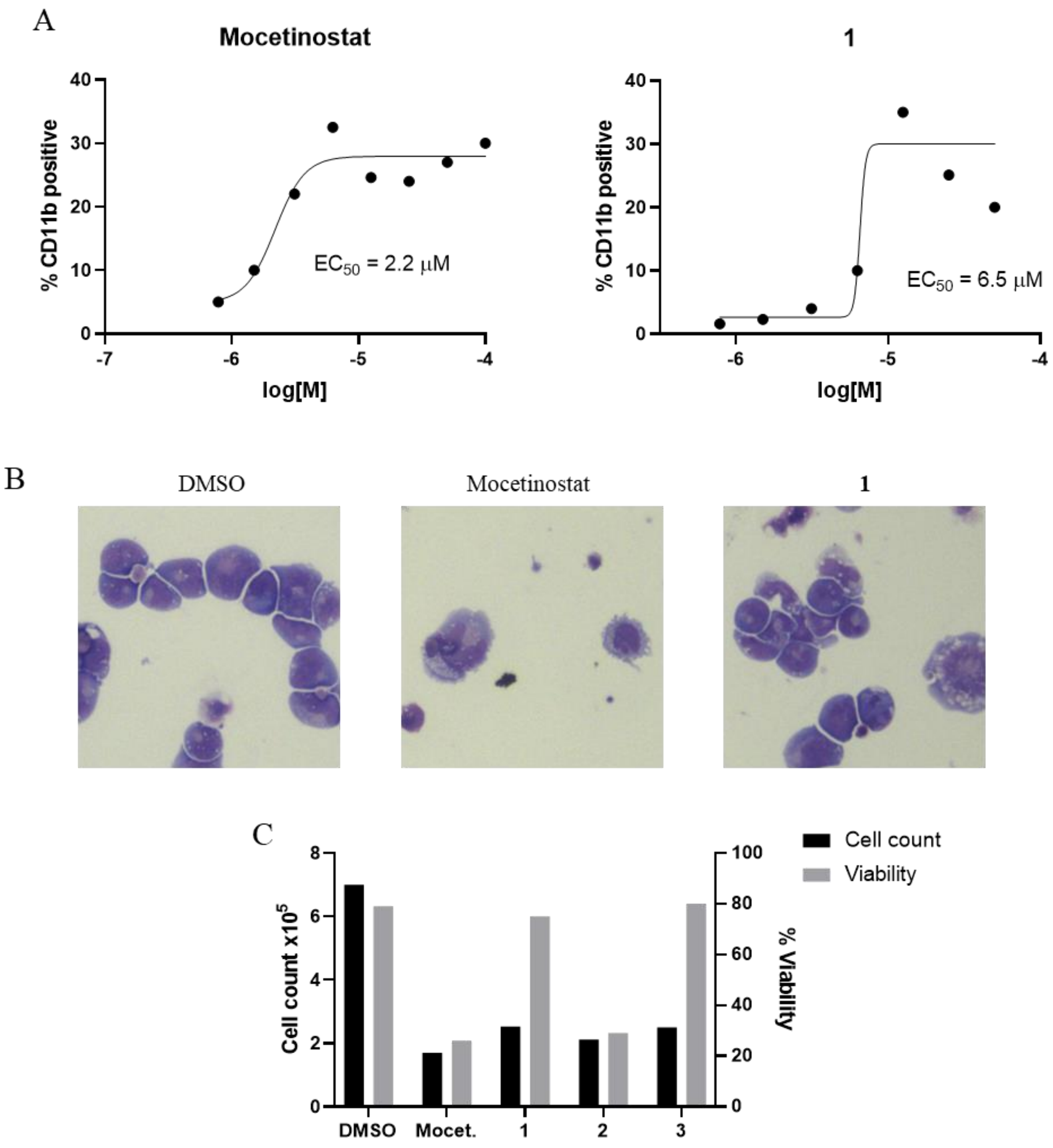


**Figure S2**: Mocetinostat and **1** induce differentiation in OCI-AML3 cells. (A) Concentration-response curves of %CD11b expression in OCI-AML3 cells treated with mocetinostat and **1**, measured by flow cytometry. (B) Cytospins of OCI-AML3 cells stained with Wright-Giemsa showed morphologic changes consistent with differentiation when treated with mocetinostat (600 nM) or **1** (2.5 μM) compared to DMSO control cells. (C) Live cells per well and cell viability measured via acridine orange and PI staining of OCI-AML3 cells treated with the compounds (600 nM mocetinostat, 10 μM **1**, 80 μM **2** and **3**) for 4 days.

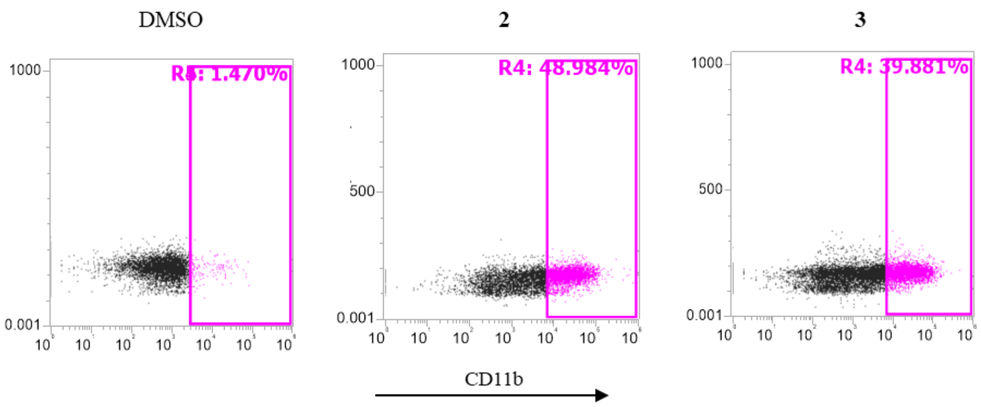


**Figure S3**. Flow cytometry plots of CD11b expression of HL60 cells treated with either DMSO control, **2** (40 μM), or **3** (40 μM).

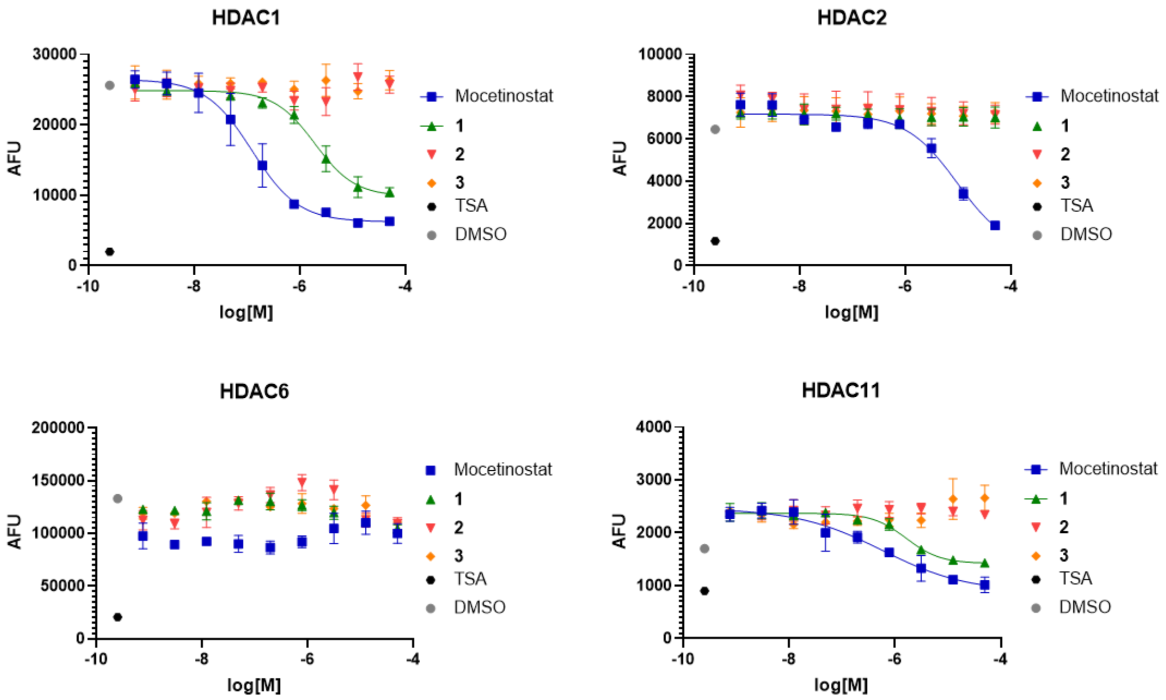


**Figure S4**: Dose-response curves of compounds in HDAC1, 2, 6, and 11 recombinant assays. The controls included DMSO as vehicle control and TSA (500 nM) as the positive control. Results are shown as means based on experiments performed in biological triplicate. Resulting $IC_{50}$ values are shown in Table 1 of the main text. AFU = Arbitraty Fluorescence Units; TSA = trichostatin

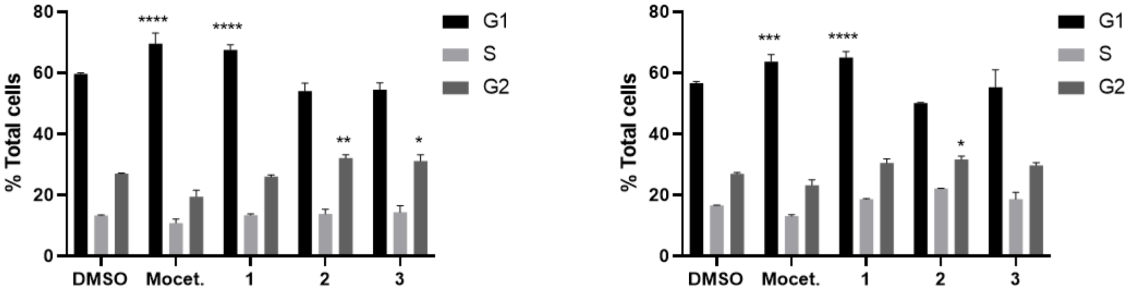


**Figure S5**. Effect of compounds on cell cycle. (A) THP1 and (B) OCI-AML3 cells were treated with the compounds (600 nM mocetinostat, 2.5 μM **1**, 30 μM **2** and **3**) for 48 h, stained with PI, and analysed by flow cytometry. Error bars shown as SEM of n=3. P values were calculated using two-way ANOVA, ****P< 0.0001, ***P< 0.001, **P<0.01, *P<0.05.

A

| Index | Name | P-value | Adjusted p-value | Odds ratio | Combined score |
|---|---|---|---|---|---|
| **1** | MACROPHAGE | 2.945E-64 | 3.181E-62 | 1.66 | 242.74 |
| **2** | NEUTROPHIL | 9.864E-30 | 5.327E-28 | 1.43 | 95.75 |
| **3** | PERIPHERAL BLOOD | 1.047E-20 | 3.771E-19 | 1.35 | 62.31 |
| **4** | FIBROBLAST | 1.302E-11 | 3.515E-10 | 1.25 | 31.42 |
| **5** | PLASMACYTOID DENDRITIC CELL | 4.623E-09 | 9.985E-08 | 1.22 | 23.37 |
| **6** | GRANULOCYTE | 8.448E-09 | 1.521E-07 | 1.21 | 22.56 |
| **7** | BLOOD PBMC | 2.049E-08 | 3.161E-07 | 1.21 | 21.38 |
| **8** | NATURAL KILLER CELLS | 3.338E-07 | 4.506E-06 | 1.19 | 17.72 |
| **9** | ALVEOLAR MACROPHAGE | 5.661E-07 | 6.793E-06 | 1.18 | 17.03 |
| **10** | SPLEEN (BULK TISSUE) | 2.186E-04 | 2.361E-03 | 1.13 | 9.55 |

B

| Index | Name | P-value | Adjusted p-value | Odds ratio | Combined score |
|---|---|---|---|---|---|
| **1** | LJP006_HME1_24H-HG-6-64-01-0.04 | 1.592E-36 | 5.275E-32 | 9.88 | 814.15 |
| **2** | LJP006_HA1E_24H-HG-6-64-01-0.04 | 5.068E-27 | 1.199E-23 | 11.67 | 706.87 |
| **3** | LJP006_HME1_24H-KIN001-043-1.11 | 1.723E-32 | 1.903E-28 | 9.17 | 670.34 |
| **4** | LJP008_PC3_24H-CGK-733-10 | 1.752E-34 | 2.902E-30 | 8.24 | 640.28 |
| **5** | LJP006_HCC515_24H-KIN001-043-10 | 2.318E-27 | 6.982E-24 | 9.71 | 595.52 |
| **6** | LJP005_A375_24H-erlotinib-10 | 2.676E-29 | 1.266E-25 | 8.99 | 591.65 |
| **7** | LJP006_HME1_3H-PD-173074-10 | 4.539E-25 | 8.352E-22 | 10.49 | 587.85 |
| **8** | LJP005_A375_24H-pelitinib-0.37 | 8.420E-29 | 3.486E-25 | 9.07 | 586.50 |
| **9** | LJP005_PC3_24H-pelitinib-3.33 | 1.396E-21 | 1.101E-18 | 11.98 | 575.10 |
| **10** | LJP006_HME1_24H-HG-6-64-01-0.12 | 3.228E-30 | 2.673E-26 | 8.32 | 564.75 |

**Figure S6**. (A) EnrichR analysis of mocetinostat, (B) compared to gene expression changes of other molecules in the L1000CDS2 database.

A

| Index | Name | P-value | Adjusted p-value | Odds ratio | Combined score |
|---|---|---|---|---|---|
| **1** | MACROPHAGE | 3.303E-68 | 3.567E-66 | 1.64 | 254.54 |
| **2** | NEUTROPHIL | 2.767E-22 | 1.494E-20 | 1.35 | 66.87 |
| **3** | PERIPHERAL BLOOD | 1.607E-18 | 5.787E-17 | 1.31 | 53.78 |
| **4** | FIBROBLAST | 2.755E-15 | 7.438E-14 | 1.28 | 42.92 |
| **5** | BLOOD PBMC | 6.674E-08 | 1.442E-06 | 1.19 | 19.63 |
| **6** | LUNG (BULK TISSUE) | 6.674E-08 | 1.201E-06 | 1.19 | 19.63 |
| **7** | SPLEEN (BULK TISSUE) | 4.271E-07 | 6.590E-06 | 1.18 | 17.24 |
| **8** | ALVEOLAR MACROPHAGE | 1.912E-06 | 2.581E-05 | 1.16 | 15.33 |
| **9** | PLASMACYTOID DENDRITIC CELL | 2.434E-06 | 2.921E-05 | 1.16 | 15.03 |
| **10** | ASTROCYTE | 3.917E-06 | 4.230E-05 | 1.16 | 14.43 |

B

| Index | Name | P-value | Adjusted p-value | Odds ratio | Combined score |
|---|---|---|---|---|---|
| **1** | LJP008_A549_24H-entinostat-10 | 8.133E-26 | 1.347E-21 | 2.63 | 151.77 |
| **2** | LJP007_HT29_24H-vorinostat-10 | 7.475E-26 | 2.476E-21 | 2.51 | 145.37 |
| **3** | LJP007_MCF7_24H-vorinostat-10 | 1.097E-25 | 1.211E-21 | 2.47 | 142.16 |
| **4** | LJP008_A549_24H-pracinostat-3.33 | 1.708E-24 | 1.131E-20 | 2.47 | 135.03 |
| **5** | LJP009_MCF7_24H-vorinostat_10 | 1.224E-24 | 1.014E-20 | 2.40 | 132.41 |
| **6** | LJP008_MCF7_24H-vorinostat-10 | 4.370E-22 | 1.316E-18 | 2.68 | 131.94 |
| **7** | LJP008_A549_24H-mocetinostat-10 | 3.609E-23 | 1.708E-19 | 2.44 | 126.23 |
| **8** | LJP008_A549_24H-pracinostat-1.11 | 2.207E-21 | 6.092E-18 | 2.54 | 120.59 |
| **9** | LJP008_MCF7_24H-belinostat-10 | 3.325E-23 | 1.836E-19 | 2.32 | 120.07 |
| **10** | LJP005_HT29_24H-XMD16-144-10 | 7.510E-23 | 2.764E-19 | 2.36 | 120.07 |

**Figure S7**. (A) EnrichR analysis of **1**, (B) compared to gene expression changes of other molecules in the L1000CDS2 database.

| Index | Name | P-value | Adjusted p-value | Odds ratio | Combined score |
|---|---|---|---|---|---|
| **1** | LJP006_HME1_24H-HG-6-64-01-0.04 | 1.592E-36 | 5.275E-32 | 9.88 | 814.15 |
| **2** | LJP006_HA1E_24H-HG-6-64-01-0.04 | 5.068E-27 | 1.199E-23 | 11.67 | 706.87 |
| **3** | LJP006_HME1_24H-KIN001-043-1.11 | 1.723E-32 | 1.903E-28 | 9.17 | 670.34 |
| **4** | LJP008 PC3_24H-CGK-733-10 | 1.752E-34 | 2.902E-30 | 8.24 | 640.28 |
| **5** | LJP006 HCC515_24H-KIN001-043-10 | 2.318E-27 | 6.982E-24 | 9.71 | 595.52 |
| **6** | LJP005_A375_24H-erlotinib-10 | 2.676E-29 | 1.266E-25 | 8.99 | 591.65 |
| **7** | LJP006_HME1_3H-PD-173074-10 | 4.539E-25 | 8.352E-22 | 10.49 | 587.85 |
| **8** | LJP005_A375_24H-pelitinib-0.37 | 8.420E-29 | 3.486E-25 | 9.07 | 586.50 |
| **9** | LJP005 PC3_24H-pelitinib-3.33 | 1.396E-21 | 1.101E-18 | 11.98 | 575.10 |
| **10** | LJP006_HME1_24H-HG-6-64-01-0.12 | 3.228E-30 | 2.673E-26 | 8.32 | 564.75 |

**Figure S8**. EnrichR analysis of **2** and **3** compared to gene expression changes of other molecules in the L1000CDS2 database.

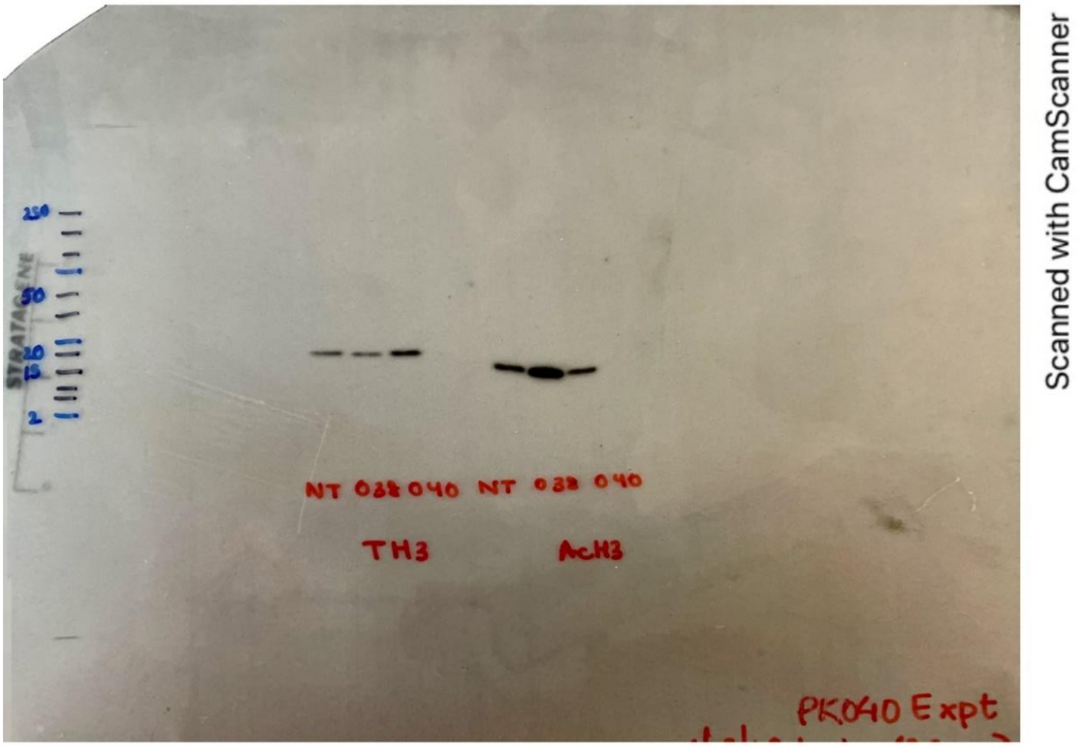


**Figure S9**. Western blot showing the effect of **1** (2.5 µM) and its *meta* analogue **3** (30 µM) on histone H3 acetylation after 24 h treatment in HL60 cells.

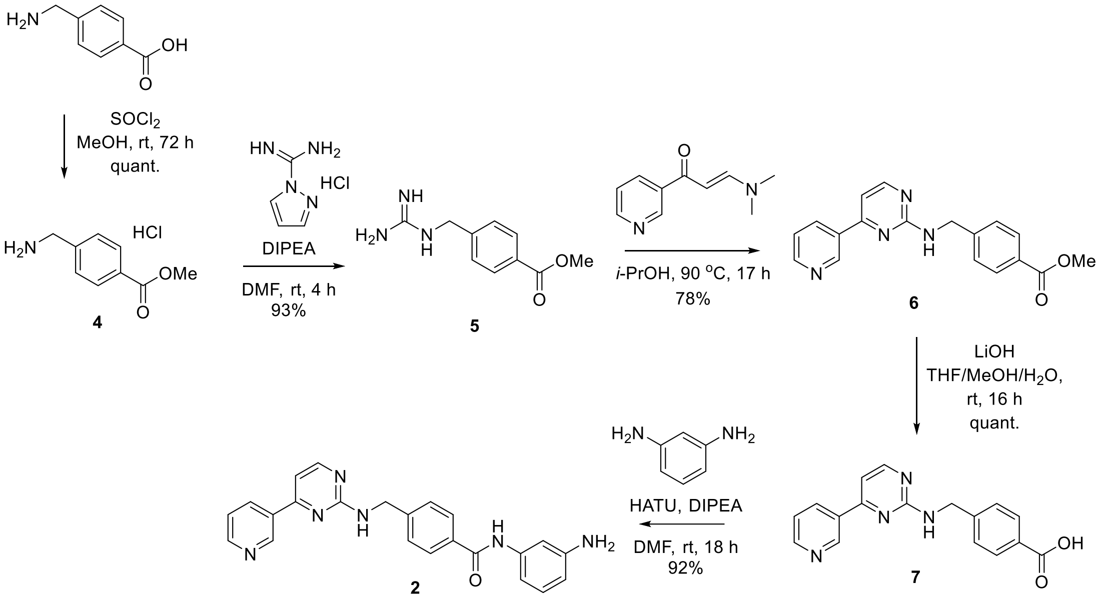


**Scheme S1**. Synthesis of *meta* analogue **2**.[1]

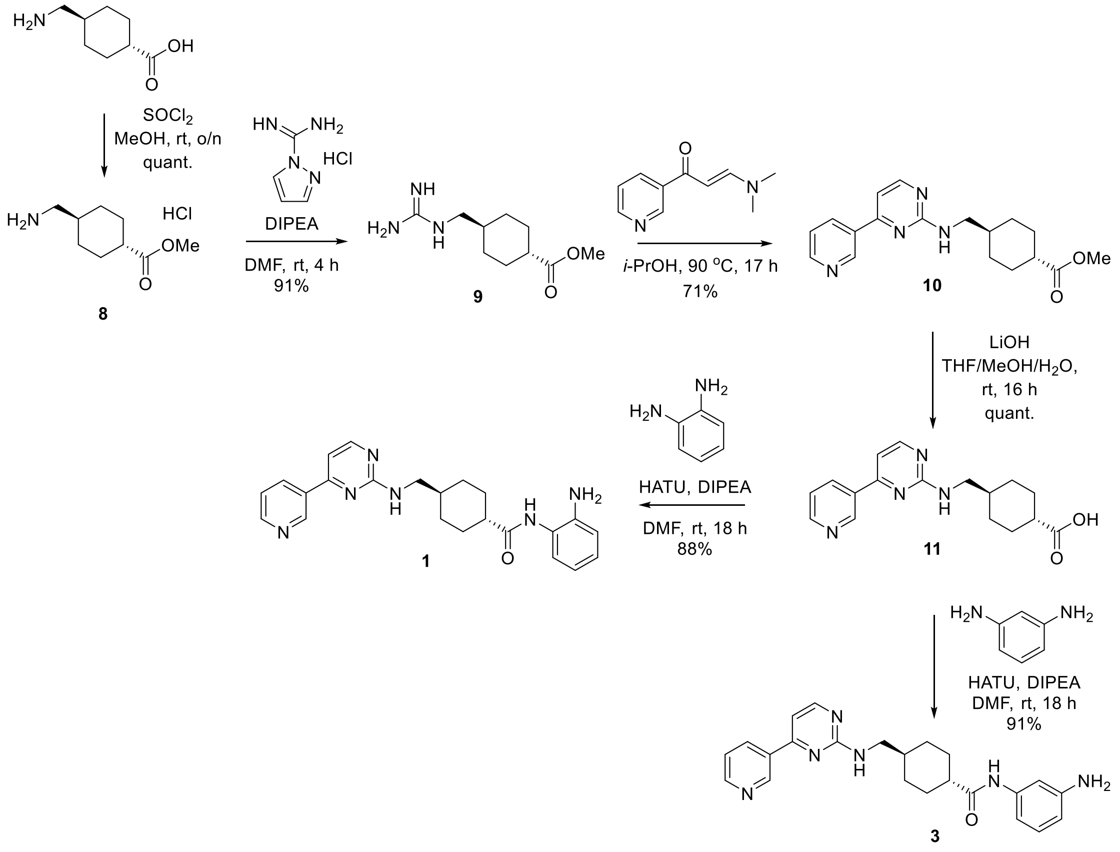


**Scheme S2**. Synthesis of **1** and its *meta* analogue **3**.

### Chemistry Experimental Procedures

#### General Information

All reactions involving moisture-sensitive reagents were carried out under nitrogen atmosphere using standard vacuum line techniques and glassware was dried and cooled before use. Anhydrous solvents were by passing over an activated alumina column, under an inert atmosphere, using a solvent purification system. Water was purified by an Elix® UV-10 system. All other reagents were used as supplied (analytical or HPLC grade) without prior purification.

Analytical thin layer chromatography (TLC) was carried out on Merck Kieselgel 60 F254 plates which were visualised using UV light (254 nm) and staining with 1% aq $KMnO_4$ followed by heating. *Flash* column chromatography was performed on Kieselgel 60 silica gel on a glass column.

Melting points (m.p.) were recorded on an EZ-Melt apparatus and are uncorrected. IR spectra were recorded on a Bruker Tensor 27 FT-IR spectrometer as thin film samples. Selected characteristic peaks are reported in wavenumbers ($cm^{-1}$). NMR spectra were recorded on Bruker AV400 or AV500 spectrometers in the deuterated solvent stated. The field was locked by external referencing to the relevant deuteron resonance. Spectra were recorded at room temperature. Chemical shifts (δ) are reported in parts per million (ppm). The multiplicity of each signal is indicated by: s (singlet); br. s (broad singlet); d (doublet); t (triplet); q (quartet); m (multiplet). Coupling constants (*J*) are quoted in Hz.

Low resolution mass spectra were recorded on an Agilent 1260 Infinity II with Diode Array and Single Quadrupole Detectors. *m/z* values are reported in Daltons and followed by their percentage abundance in parentheses. Accurate mass measurements were run on Bruker Micro TOF spectrometer internally calibrated with polyalanine.

Mocetinostat was purchased from Key Organics (MFCD10565970).

##### Methyl 4-(aminomethyl)benzoate hydrochloride (4)

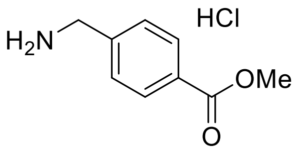


To 4-aminobenzoic acid (6.00 g, 39.7 mmol) in MeOH (400 mL) was added thionyl chloride (6.5 mL, 90 mmol) at 0 °C. The mixture was stirred at room temperature for 72 h and concentrated *in vacuo* to afford methyl ester **4** (quant.) that was used in the next step without further purification.

##### Methyl 4-(guanidinomethyl)benzoate (5)[2]

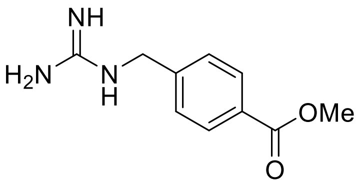

A mixture of amine **4** (500 mg, 3.02 mmol) and DIPEA (1.16 mL, 6.64 mmol) in DMF (5.0 mL) was stirred for 10 min before addition of 1*H*-pyrazole-1-carboxamidine hydrochloride (487 mg, 3.32 mmol). The resulting solution was stirred until clear (4 h); it was concentrated *in vacuo* before addition of a saturated aqueous solution of $NaHCO_3$ to give a suspension, which was filtered and washed with cold water to afford guanidine **5** (585 mg, 2.82 mmol, 93%) as a white solid. $^1$H NMR (400 MHz, DMSO-$d_6$) δ 7.95 (d, *J* = 8.3 Hz, 2H), 7.42 (d, *J* = 8.3 Hz, 2H), 4.41 (s, 2H), 3.85 (s, 3H); *m/z* (ESI$^+$) 208 ([M+H]$^+$).

**Methyl 4-(((4-(pyridin-3-yl)pyrimidin-2-yl)amino)methyl)benzoate (6)**[2]

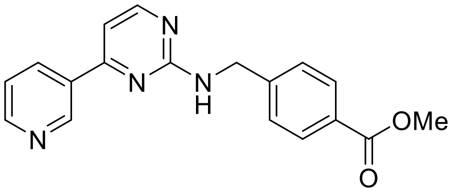


Guanidine **5** (260 mg, 1.25 mmol) was dissolved in *i*-PrOH (4.0 mL) before addition of molecular sieves (4 Å) followed by (*E*)-3-(dimethylamino)-1-(pyridin-3-yl)prop-2-en-1-one (265 mg, 1.51 mmol). The suspension was heated for 90 °C for 16 h, the temperature lowered to 70 °C before addition of MeOH (1.0 mL), and the reaction heated to 80 °C for 30 min. The suspension was cooled to room temperature and filtered through Celite™ using MeOH as an eluent. The filtrate was concentrated *in vacuo* and triturated ($Et_2O$/pentane) to yield title compound **6** (312 mg, 0.974 mmol, 78%) as an off-white solid; m.p. 175 ºC (lit m.p. 174-175 °C[3]). $^1$H NMR ($CDCl_3$, 400 MHz) δ 9.19 (dd, *J* = 2.2, 0.9 Hz, 1H), 8.69 (dd, *J* = 4.8, 2.0 Hz, 1H), 8.40 (d, *J* = 5.2 Hz, 1H), 8.27 (dt, *J* = 8.0, 2.0 Hz, 1H), 8.05 – 7.97 (m, 2H), 7.46 (d, *J* = 8.3 Hz, 2H), 7.39 (ddd, *J* = 8.0, 4.8, 0.9 Hz, 1H), 7.04 (d, *J* = 5.2 Hz, 1H), 5.67 (t, *J* = 6.2 Hz, 1H), 4.80 (d, *J* = 6.2 Hz, 2H), 3.90 (s, 3H); HRMS (ESI$^+$) calculated for $C_{18}H_{17}N_4O_2$ ([M+H]$^+$) 321.1346, found 321.1347.

**4-(((4-(Pyridin-3-yl)pyrimidin-2-yl)amino)methyl)benzoic acid (7)**[2]

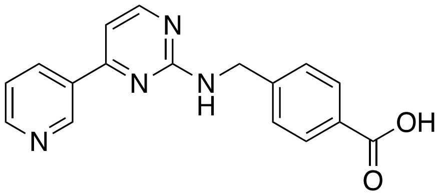


Methyl ester **6** (126 mg, 0.394 mmol) was dissolved in THF/MeOH (4:1, 5.0 mL) before addition of 1 M aqueous LiOH until pH>8. The resulting reaction was stirred for 16 h at room temperature, after which it was acidified with 1 M aqueous HCl until pH<5. The solution was concentrated *in vacuo* and the obtained carboxylic acid **8** (quant.) was used in the next step without further purification; m.p. 223 °C (lit m.p. 223-224 °C[3]). $^1$H NMR (500 MHz, DMSO-$d_6$) δ 9.36 (br s, 1H), 8.90 – 8.78 (m, 2H), 8.48 (br s, 1H), 8.33 (br s, 1H), 7.89 (d, *J* = 7.9 Hz, 2H), 7.56 – 7.31 (m, 4H), 4.66 (br s, 2H); HRMS (ESI$^+$) calculated for $C_{17}H_{15}N_4O_2$ ([M+H]$^+$) 307.1190, found 307.1189. NMR data are in agreement with the reported values.[4]

**N-(3-Aminophenyl)-4-(((4-(pyridin-3-yl)pyrimidin-2-yl)amino)methyl)benzamide (2)**

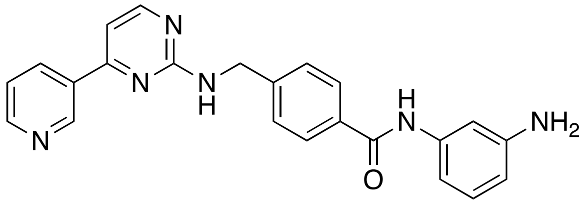

Acid **7** (119 mg, 0.380 mmol) was dissolved in DMF (3.0 mL) before sequential addition of DIPEA (273 µL, 1.56 mmol), 1,3-phenylenediamine (51 mg, 0.47 mmol) and HATU (207 mg, 0.546 mmol). The resulting solution was stirred for 18 h, diluted with EtOAc and washed with 3xbrine/water (1:1). The organic phase was dried over anhydrous $Na_2SO_4$, filtered, and concentrated *in vacuo*. The crude material was purified on silica gel (4% MeOH in $CH_2Cl_2$) to give benzamide **2** (139 mg, 0.351 mmol, 92%) as a white solid; m.p. 191 ºC. $^1$H NMR (400 MHz, DMSO-$d_6$) 9.85 (s, 1H), 9.24 (s, 1H), 8.68 (d, *J* = 3.3 Hz, 1H), 8.44 – 8.36 (m, 2H), 7.98 (t, *J* = 6.3 Hz, 1H), 7.86 (d, *J* = 8.3 Hz, 2H), 7.56 – 7.45 (m, 3H), 7.25 (d, *J* = 5.1 Hz, 1H), 7.07 (t, *J* = 2.1 Hz, 1H), 6.94 (t, *J* = 7.9 Hz, 1H), 6.83 (d, *J* = 8.6 Hz, 1H), 6.32 – 6.26 (m, 1H), 5.05 (br s, 2H), 4.65 (d, *J* = 6.3 Hz, 2H); $^{13}$C NMR (126 MHz, DMSO-$d_6$) δ 165.2, 162.4, 161.0, 159.4, 151.3, 148.9, 148.0, 144.1, 139.8, 134.2, 133.6, 132.4, 128.8, 127.6, 126.9, 123.8, 109.7, 108.4, 106.2, 106.1, 44.0; HRMS (ESI$^+$) calculated for $C_{23}H_{21}N_6O$ ([M+H]$^+$) 397.1717, found 397.1790.

**Methyl *trans*-4-(aminomethyl)cyclohexane-1-carboxylate hydrochloride (8)**

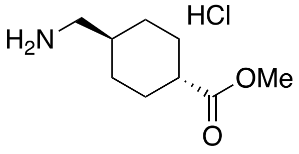

To a suspension of *trans*-4-(aminomethyl)cyclohexane-1-carboxylic acid (6.0 g, 38 mmol) in MeOH (95 mL) was added thionyl chloride (3.8 mL, 52.4 mmol) at 0 °C. The mixture was stirred at room temperature overnight and then concentrated *in vacuum* to give methyl ester **8** (7.89 g, 38.0 mmol, quant.) as a light green solid; m.p. 179 ºC (lit m.p. 168-170 °C[5]). $\nu_{max}$/cm$^{-1}$ 2938, 1723, 1173; $^1$H NMR (400 MHz, DMSO-$d_6$) δ 3.49 (s, 3H), 2.53 (t, *J* = 6.2 Hz, 2H), 2.41 (s, 2H), 2.15 (tt, *J* = 12.1, 3.5 Hz, 1H), 1.86 – 1.77 (m, 2H), 1.75 – 1.69 (m, 2H), 1.46 (m, 1H), 1.19 (qd, *J* = 12.9, 3.3 Hz, 2H), 0.88 (qd, *J* = 12.9, 3.4 Hz, 2H); $^{13}$C NMR (100 MHz, DMSO-$d_6$) δ 175.7, 51.8, 44.6, 42.4, 35.3, 29.1, 28.3; *m/z* (ESI$^+$) 172.1 ([M+H]$^+$, 100%); HRMS (ESI$^+$) calculated for $C_9H_{18}O_2N$ ([M+H]$^+$) 172.1332, found 172.1330.

**Methyl *trans*-4-(guanidinomethyl)cyclohexane-1-carboxylate (9)**

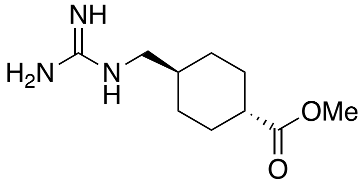

A mixture of amine **8** (500 mg, 3.02 mmol) and DIPEA (1.16 mL, 6.64 mmol) in DMF (5.0 mL) was stirred for 10 min before addition of 1*H*-pyrazole-1-carboxamidine hydrochloride (487 mg, 3.32 mmol). The resulting solution was stirred until clear (4 h); it was concentrated *in vacuo* before addition of saturated aqueous $NaHCO_3$ to give a suspension, which was filtered and washed with cold water to afford guanidine **9** (585 mg, 2.74 mmol, 91%) as a white solid. $^{1}$H NMR (400 MHz, DMSO-*d*$_6$) δ 3.58 (s, 3H), 2.90 (d, *J* = 6.5 Hz, 2H), 2.23 (tt, *J* = 12.2, 3.6 Hz, 1H), 1.90 (dd, *J* = 13.5, 3.5 Hz, 2H), 1.76 (d, *J* = 12.8 Hz, 2H), 1.48 – 1.38 (m, 1H), 1.29 (qd, *J* = 13.0, 3.3 Hz, 2H), 0.93 (q, *J* = 14.2, 13.1 Hz, 2H); *m/z* (ESI$^+$) 214.2 ([M+H]$^+$); HRMS (ESI$^+$) calculated for $C_{10}H_{20}N_3O_2$ ([M+H]$^+$) 214.1550, found 214.1549.

**Methyl *trans*-4-(((4-(pyridin-3-yl)pyrimidin-2-yl)amino)methyl)cyclohexane-1-carboxylate (10)**

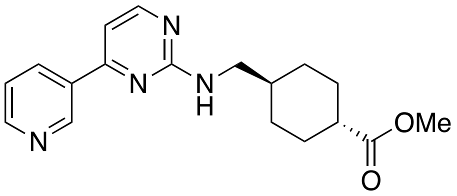


Guanidine **9** (180 mg, 0.840 mmol) was dissolved in *i*-PrOH (3.0 mL) before addition of molecular sieves (4 Å) followed by (*E*)-3-(dimethylamino)-1-(pyridin-3-yl)prop-2-en-1-one (180 mg, 1.01 mmol). The suspension was heated for 90 °C for 16 h, the temperature lowered to 70 °C before addition of MeOH (1 mL), and the reaction heated to 80 °C for 30 min. The suspension was cooled to room temperature and filtered through Celite™ using MeOH as an eluent. The filtrate was concentrated *in vacuo* and triturated ($Et_2O$/pentane) to yield title compound **10** (195 mg, 0.597 mmol, 71%) as an off-white solid; m.p. 220 ºC. $^{1}$H NMR (500 MHz, MeOD-*d*$_4$) δ 9.22 (d, *J* = 2.2 Hz, 1H), 8.65 (dd, *J* = 4.9, 1.6 Hz, 1H), 8.49 (dt, *J* = 8.0, 2.0 Hz, 1H), 8.34 (d, *J* = 5.2 Hz, 1H), 7.58 (dd, *J* = 8.0, 4.9 Hz, 1H), 7.15 (d, *J* = 5.2 Hz, 1H), 3.65 (s, 3H), 3.35 – 3.32 (m, 2H), 2.30 (tt, *J* = 12.3, 3.6 Hz, 1H), 2.04 – 1.96 (m, 2H), 1.96 – 1.89 (m, 2H), 1.72 – 1.60 (m, 1H), 1.40 (qd, *J* = 13.0, 3.4 Hz, 2H), 1.08 (qd, *J* = 13.0, 3.4 Hz, 2H); $^{13}$C NMR (125 MHz, MeOD-*d*$_4$) δ 178.7, 164.0, 163.6, 160.0, 151.6, 148.7, 136.6, 134.8, 125.5, 106.8, 52.3, 48.3, 44.5, 38.5, 31.0, 29.8; *m/z* (ESI$^+$) 327.1 ([M+H]$^+$); HRMS (ESI$^+$) calculated for $C_{18}H_{23}N_4O_2$ ([M+H]$^+$) 327.1815, found 327.1810.

***trans*-4-(((4-(Pyridin-3-yl)pyrimidin-2-yl)amino)methyl)cyclohexane-1-carboxylic acid (11)**

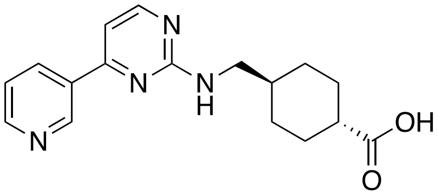


Methyl ester **10** (110 mg, 0.337 mmol) was dissolved in THF/MeOH (4:1, 5.0 mL) before addition of aqueous 1 M LiOH until pH>8. The resulting reaction was stirred for 16 h at room temperature, after which it was acidified with aqueous 1 M HCl until pH<5. The solution was concentrated *in vacuo* and the obtained carboxylic acid **11** (quant.) was used in the next step without further purification. $^{1}$H NMR (400 MHz, MeOD-*d*$_4$) δ 9.42 (s, 1H), 8.89 (d, *J* = 8.1 Hz, 1H), 8.84 (d, *J* = 5.4 Hz, 1H), 8.41 (d, *J* = 5.7 Hz, 1H), 7.89 (dd, *J* = 8.1, 5.2 Hz, 1H), 7.38 (d, *J* = 5.7 Hz, 1H), 3.36 – 3.26 (m, 2H), 2.26 (tt, *J* = 12.1, 3.5 Hz, 1H), 2.07 – 1.92 (m, 4H),

1.76 – 1.65 (m, 1H), 1.43 (qd, $J$ = 13.0, 3.4 Hz, 2H), 1.12 (q, $J$ = 12.7 Hz, 2H); HRMS (ESI$^+$) calculated for $C_{17}H_{21}N_4O_2$ ([M+H]$^+$) 313.1659, found 313.1657.

***trans-N*-(2-Aminophenyl)-4-(((4-(pyridin-3-yl)pyrimidin-2-yl)amino)methyl)cyclohexane-1-carboxamide (1)**

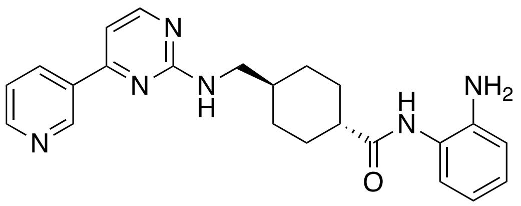


Acid **11** (640 mg, 2.06 mmol) was dissolved in DMF (6.0 mL) before sequential addition of DIPEA (1.44 mL, 8.22 mmol), *o*-phenylenediamine (269 mg, 2.47 mmol), and HATU (1.09 g, 2.88 mmol). The resulting solution was stirred for 18 h, then diluted with EtOAc and washed with 5xbrine/water (1:1). The organic phase was dried over anhydrous $Na_2SO_4$, filtered, and concentrated *in vacuo*. The crude material was purified on silica gel (4% MeOH in $CH_2Cl_2$) to give benzamide **1** (730 mg, 1.81 mmol, 88%) as a white solid; m.p. 179 ºC. $^1$H NMR (500 MHz, MeOD-$d_4$) δ 9.25 (s, 1H), 8.67 – 8.62 (m, 1H), 8.51 (dt, $J$ = 8.0, 1.9 Hz, 1H), 8.34 (d, $J$ = 5.2 Hz, 1H), 7.57 (dd, $J$ = 8.1, 4.9 Hz, 1H), 7.15 (d, $J$ = 5.2 Hz, 1H), 7.06 (dd, $J$ = 7.9, 1.5 Hz, 1H), 7.02 (td, $J$ = 7.6, 1.5 Hz, 1H), 6.84 (dd, $J$ = 8.0, 1.4 Hz, 1H), 6.71 (td, $J$ = 7.5, 1.4 Hz, 1H), 3.39 – 3.33 (m, 2H), 2.43 (tt, $J$ = 12.2, 3.3 Hz, 1H), 2.05 – 1.97 (m, 4H), 1.78 – 1.70 (m, 1H), 1.60 (qd, $J$ = 13.8, 13.2, 3.6 Hz, 2H), 1.37 (dd, $J$ = 6.9, 3.1 Hz, 2H), 1.23 – 1.10 (m, 1H); $^{13}$C NMR (125 MHz, MeOD-$d_4$) δ 178.0, 164.2, 160.0, 151.6, 148.9, 143.1, 136.5, 135.0, 128.2, 127.0, 125.4, 125.3, 119.7, 118.6, 111.7, 106.7, 46.6, 39.0, 38.7, 31.2, 30.5; *m/z* (ESI$^+$) 403.2 ([M+H]$^+$); HRMS (ESI$^+$) calculated for $C_{23}H_{27}N_6O$ ([M+H]$^+$) 403.2241, found 403.2236.

***trans-N*-(3-Aminophenyl)-4-(((4-(pyridin-3-yl)pyrimidin-2-yl)amino)methyl)cyclohexane-1-carboxamide (3)**

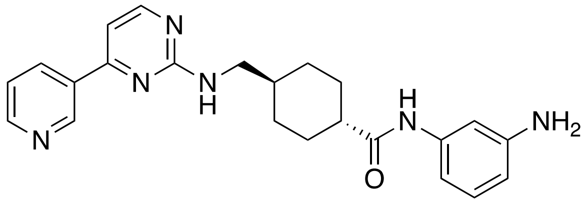


Acid **11** (122 mg, 0.390 mmol) was dissolved in DMF (3.0 mL) before sequential addition of DIPEA (273 µL, 1.56 mmol), 1,3-phenylenediamine (51 mg, 0.47 mmol), and HATU (207 mg, 0.546 mmol). The resulting solution was stirred for 18 h, then diluted with EtOAc and washed with 3xbrine/water (1:1). The organic phase was dried over anhydrous $Na_2SO_4$, filtered, and concentrated *in vacuo*. The crude material was purified on silica gel (4% MeOH in $CH_2Cl_2$) to give benzamide **3** (143 mg, 0.355 mmol, 91%) as a white solid; m.p. 180 ºC. $^1$H NMR (500 MHz, MeOD-$d_4$) δ 9.25 (s, 1H), 8.64 (dd, $J$ = 4.9, 1.6 Hz, 1H), 8.51 (dt, $J$ = 7.9, 1.8, 1H), 8.34 (d, $J$ = 5.2 Hz, 1H), 7.57 (dd, $J$ = 8.0, 4.9 Hz, 1H), 7.15 (d, $J$ = 5.2 Hz, 1H), 7.05 – 6.96 (m, 2H), 6.80 (ddd, $J$ = 8.0, 2.0, 1.0 Hz, 1H), 6.46 (ddd, $J$ = 8.0, 2.3, 1.0 Hz, 1H), 3.39 – 3.33 (m, 2H), 2.33 (tt, $J$ = 12.4, 3.2 Hz, 1H), 2.05 – 1.88 (m, 3H), 1.72 (br s, 1H), 1.63 – 1.50 (m, 2H), 1.41 – 1.25 (m, 3H), 1.21 – 1.07 (m, 2H); $^{13}$C NMR (101 MHz, MeOD-$d_4$) δ 177.4, 164.2, 160.1, 160.0, 151.6, 149.2, 148.8, 140.7, 136.5, 134.9, 130.2, 125.3, 112.5, 111.2, 108.5, 106.7,

47.2, 39.6, 38.7, 31.2, 30.3; *m/z* (ESI$^+$) 403.3 ([M+H]$^+$); HRMS (ESI$^+$) calculated for $C_{23}H_{27}N_6O$ ([M+H]$^+$) 403.2241, found 403.2240.
